# 15-PGDH inhibition promotes hematopoietic recovery and enhances HSC function during aging

**DOI:** 10.1101/2025.04.11.648417

**Authors:** Rahul Chaudhary, Brittany A. Cordova, Marcus Hong, Bailey R. Klein, Lyannah A. Contreras, Ritisha Rashmil, Filip Goshevski, Julianne N.P. Smith, Derek J. Taylor, Andrew A. Pieper, Sanford Markowitz, Amar B. Desai

**Affiliations:** Department of Medicine, and Case Comprehensive Cancer Center Case Western Reserve University, Cleveland, OH, USA; Department of Pharmacology, Case Western Reserve University, Cleveland, OH, USA; Brain Health Medicines Center, Harrington Discovery Institute, University Hospitals Cleveland Medical Center, Cleveland, OH, USA; Department of Psychiatry, Case Western Reserve University, Cleveland, OH, USA; Geriatric Psychiatry, GRECC, Louis Stokes VA Medical Center, Cleveland, OH, USA; Institute for Transformative Molecular Medicine, School of Medicine, Case Western Reserve University, Cleveland, OH, USA; Department of Neuroscience, Case Western Reserve University, Cleveland, OH, USA; Department of Pathology, Case Western Reserve University, Cleveland, OH, USA; University Hospitals Seidman Cancer Center, Cleveland OH 44106 US

**Keywords:** aging, hematopoietic stem cells, hematopoietic stem cell transplantation, stem cells, drug target

## Abstract

Hematopoietic aging is characterized by diminished stem cell regenerative capacity and an increased risk of hematologic dysfunction. We previously identified that the prostaglandin-degrading enzyme 15-hydroxyprostaglandin dehydrogenase (15-PGDH) regulates hematopoietic stem cell activity. Here, we expand on this work and demonstrate that in aged mice, (1) 15-PGDH expression and activity remain conserved in the bone marrow and spleen, suggesting it remains a viable therapeutic target in aging, (2) prolonged PGDH inhibition (PGDHi) significantly increases the frequency and number of phenotypic hematopoietic stem and progenitor cells across multiple compartments, with transcriptional changes indicative of enhanced function, (3) PGDHi-treated bone marrow enhances short-term hematopoietic recovery following transplantation, leading to improved peripheral blood output and accelerated multilineage reconstitution, and (4) PGDHi confers a competitive advantage in primary hematopoietic transplantation while mitigating age-associated myeloid bias in secondary transplants. Notably, these effects occur without perturbing steady-state blood production, suggesting that PGDHi enhances hematopoiesis under regenerative conditions while maintaining homeostasis. Our work identifies PGDHi as a translatable intervention to rejuvenate aged HSCs and mitigate hematopoietic decline.

**Significance Statement:** We identify 15-hydroxyprostaglandin dehydrogenase inhibition (PGDHi) as a strategy to enhance hematopoietic stem cell function in aging. In aged mice, PGDHi expands stem and progenitor populations, accelerates hematopoietic recovery after transplantation, and reduces myeloid bias while maintaining steady-state blood production. These findings highlight a potential therapeutic approach to restore hematopoietic resilience and improve regenerative outcomes in aging.

**Graphical Abstract:** 15-prostaglandin dehydrogenase inhibition ameliorates multiple facets of age-related hematopoietic decline. Schematic made using BioRender.com.

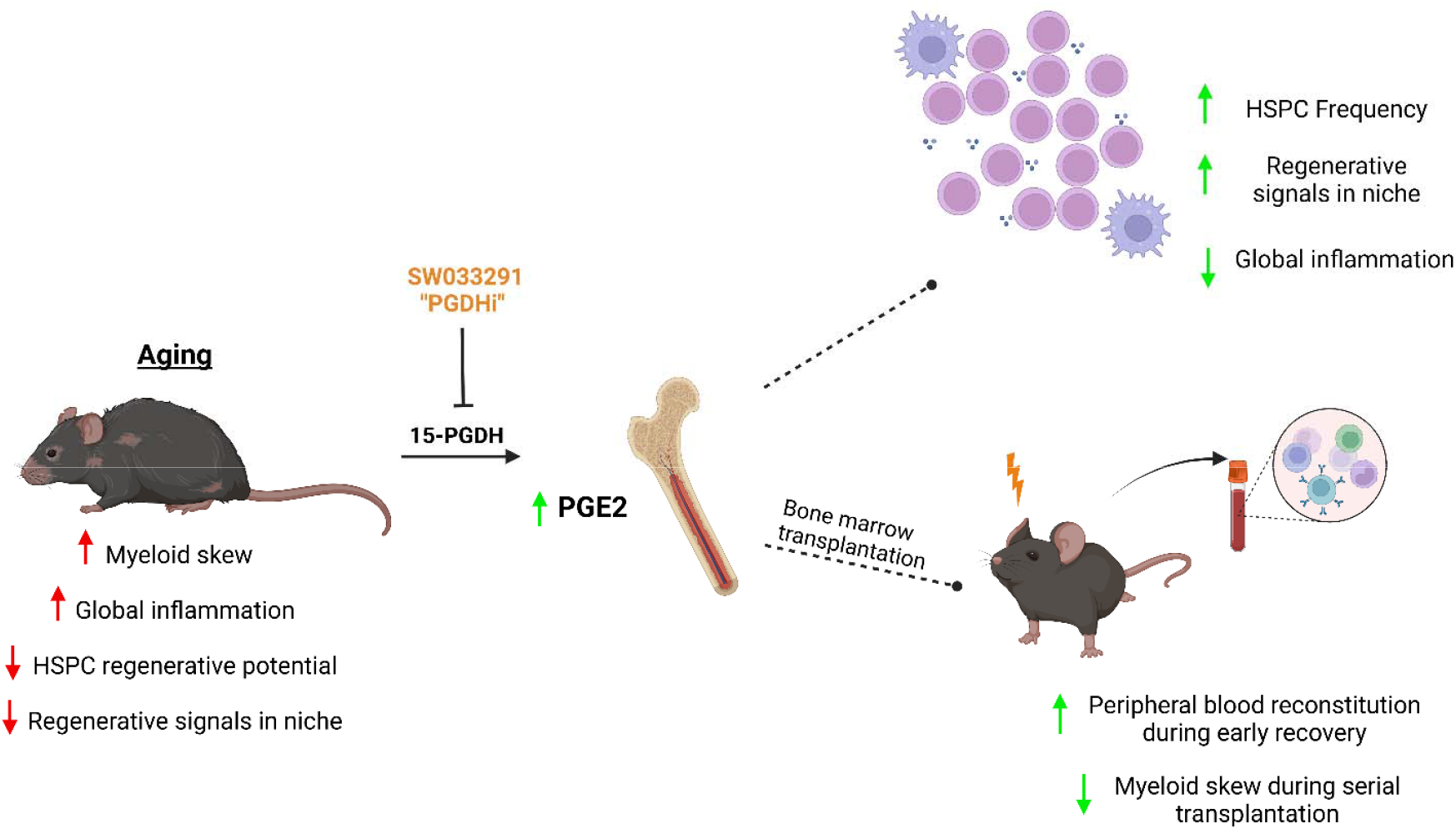

## Introduction

Aging is characterized by progressive physiologic decline, reduced tissue homeostasis, and heightened disease susceptibility. At the cellular level, this process is driven by interconnected factors, including genomic instability, epigenetic dysregulation, mitochondrial dysfunction, cellular senescence, chronic inflammation, and disrupted intercellular communication ^1 2^. These hallmarks of aging collectively impair regenerative tissues, including the hematopoietic system, which is responsible for the continuous replenishment of blood and immune cells.

Hematopoietic stem cells (HSCs) are particularly vulnerable to age-related dysfunction, leading to a decline in regenerative capacity and hematopoietic output. Aged HSCs exhibit impaired self-renewal, increased proliferation at the expense of quiescence, and a differentiation bias toward the myeloid lineage at the expense of lymphoid cells. This imbalance contributes to immunosenescence, increased susceptibility to infections, anemia, and a higher incidence of hematologic malignancies ^3 4^. Additionally, aged HSCs accumulate DNA damage due to defective DNA repair and oxidative stress, further impairing their function ^5 6^. These defects are exacerbated by inflammatory changes in the bone marrow niche, where increased levels of pro-inflammatory cytokines create a microenvironment that further drives HSC dysfunction. Chronic exposure to inflammatory signaling pathways, such as those mediated by IL-6 and TNF-α, promotes myeloid-biased differentiation while suppressing normal stem cell renewal. This persistent, low-grade inflammatory state (referred to as “inflammaging”) is a driver of hematopoietic dysfunction ^7,8^.

As a consequence of these intrinsic and microenvironmental changes, aging is linked to a rising incidence of hematologic disorders. Myelodysplastic syndromes and acute myeloid leukemia are significantly more prevalent in older individuals, in part due to the accumulation of HSC mutations and clonal hematopoiesis of indeterminate potential, which can confer a competitive advantage to pre-malignant clones ^9 10 11^. Additionally, the reduced regenerative capacity of aged HSCs exacerbates cytopenias, increasing the risk of anemia and impaired immune surveillance, which further contributes to disease susceptibility ^12,13^. These deficits also impair the response to hematologic stress, such as chemotherapy-induced neutropenia, where older patients experience delayed hematologic recovery and an increased risk of infection-related complications ^14^.

Beyond these age-associated disease risks, aged HSCs also exhibit functional deficits in transplantation. In murine competitive transplantation models, aged HSCs show reduced engraftment capacity, delayed multilineage recovery, and a persistent myeloid bias, even when transplanted into young, supportive BM niches^15,16^. These defects are clinically relevant, as older patients undergoing hematopoietic stem cell transplantation frequently experience prolonged cytopenias, poor graft function, and increased transplant failure, leading to worsened patient outcomes^17,18^. Furthermore, the aged BM microenvironment contributes to these poor outcomes by failing to support efficient HSC expansion, limiting the effectiveness of transplantation-based therapies^19^. Despite these recognized challenges, therapeutic strategies aimed at rejuvenating aged HSC function and improving transplantation outcomes remain limited.

Prostaglandin E2 (PGE2) has been demonstrated to regulate HSC function by influencing self-renewal, proliferation, and survival^20-22^. Our previous work has identified 15-hydroxyprostaglandin dehydrogenase (15-PGDH), the enzyme that degrades PGE2, as a novel regulator of HSC function. We demonstrated that inhibiting 15-PGDH (PGDHi) with the small molecule (+)SW033291 induces expression of pro-hematopoietic cytokines CXCL12 and SCF, enhances hematopoietic recovery after bone marrow transplantation in both young and aged settings, protects against immune-mediated bone marrow failure, and that aged 15-PGDH knockout mice show improved hematopoietic function^23-26^. Together this suggests that PGDHi represents a promising approach to counteract hematopoietic aging by improving HSC function, mitigating myeloid bias, and enhancing engraftment potential. In this study we examined the potential of PGDHi to prevent age-associated hematopoietic decline.

## Materials and Methods

### Study approval

Animals were housed in the AAALAC accredited facilities of the Case Western Reserve University School of Medicine. Husbandry and experimental procedures were approved by the CWRU Institutional Animal Care and Use Committee (IACUC) in accordance with approved IACUC protocol 2019-0065.

### Animals

Steady-state and transplantation analyses were performed on 8wk old female C57BL/6J mice obtained from Jackson Laboratories. B6.SJL-Ptprca Pepc/BoyJ mice for competitive transplant studies were also obtained from Jackson Laboratories. All animals were observed daily for signs of illness or distress. Mice were housed in standard microisolator cages and maintained on a defined, irradiated diet and autoclaved water.

### (+)SW033291 for PGDHi

The 15-PGDH inhibitor (+)SW033291 has been previously described ^23,24^, and was provided by Dr. Sanford Markowitz. (+)SW033291 was prepared in a vehicle of 10% ethanol, 5% Cremophor EL, 85% dextrose-5 water, at 250ug/200ul for use at 10mg/kg for a 25g mouse, and administered by intraperitoneal (I.P.) injection, twice per day, 6-8 hours apart.

### 15-PGDH activity assay

Splenic lysates were prepared using the Precellys 24 homogenizer, in a lysis buffer containing 50mM Tris HCl, 0.1mM DTT, and 0.1mM EDTA. Bone marrow was flushed, pelleted, and lysed using the same buffer, with sonication. Enzymatic activity was measured by following the transfer of tritium from a tritiated PGE2 substrate to glutamate by coupling 15-PGDH to glutamate dehydrogenase ^27^. Activity was expressed as counts per minute, per mg total protein assayed.

### Peripheral blood, bone marrow, and spleen collection

Peripheral blood (PB) was collected by puncturing the submandibular vein with a 5 mm lancet, and four drops of blood were collected into a K2 EDTA blood collection tube (Beckton Dickinson Catalog # 365974). Blood counts were analyzed using a Hemavet 950 FS hematology analyzer. For cytometric analysis total blood was lysed twice with 1x RBC Lysis Buffer (EBioScience, Catalog #00-4300-54) and the remaining white cell pellet was used for downstream cytometric analysis. Mice were sacrificed by cervical dislocation. Spleens were extracted and a single cell suspension was obtained by using the rubber end of the plunger of a 3mL syringe to mince the spleen through a 40-um cell strainer into a cell culture dish containing 1 ml of PBS + 2% FBS (FACS) buffer. For BM collection, the epiphyses were removed from the ends of both the tibias and the femurs of the mouse, and bone marrow was flushed from the medullary cavity with FACS using a 1 ml syringe fitted with a 26-gauge needle. Splenic and bone marrow cells were aliquoted and pelleted for downstream use.

### RNA extraction and quantitative PCR

Bone marrow and splenic lysates were obtained as described above. RNA was isolated using the RNeasy Mini Kit (Qiagen, Catalog # 74106) and cDNA was synthesized using a PrimeScript RT Reagent Kit (Takara, Catalog # RR037B), both according to manufacturer’s protocol. Real-time PCR (RT-PCR) was performed in a 20 μL reaction containing 1 μL cDNA template and a 1:20 dilution of primer/probe with qMAX Probe Low Rox qPCR Mix (Accuris, Catalog # PR2001-L-100). Samples were run on a Bio-Rad CFX96 optical module thermal cycler. Thermal cycling conditions were 95°C for 3 minutes, followed by 50 cycles at 95°C for 15 seconds and 60°C for 1 minute. Murine probe/primer sets for genes assayed were obtained from Life Technologies, Thermo Fisher Scientific, and were as follows: Actb Mm02619580_g1, Arg1 Mm00475988_m1, Il10 Mm01288386_m1, Mrc1 Mm01329359_m1, Tgfb1 Mm01178820_m1, HPGD Mm00515121_m1, Angpt1 Mm00456503_m1, CD86 Mm00444540, Csf1 Mm00432686_m1, Kitl Mm00442972_m1, Mrc1 Mm01329359_m1, Spp1 Mm00436767_m1, Nos2 Mm00440502_m1, Nr4a1 Mm01300401_m1, Cebpb Mm07294206_s1, Ptger2 Mm00436051_m1 and Ptger4 Mm00436053_m1. Each sample was run in triplicate and Cq values were determined as the average values the three independent RT-PCR reactions.

### Immunohistochemistry

Bone marrow plugs flushed from the tibia and total spleens were fixed in 10% NBF for 24 hours and then transferred to 70% ethanol. Samples were shipped to HistoWiz, where they were embedded in paraffin and sectioned at 4 μm. IHC was performed according to HistoWiz protocols (https://home.histowiz.com/faq/). HistoWiz defines their standard methods as the use of a Bond Rx autostainer (Leica Biosystems) with enzyme treatment using standard protocols, and detection via Bond Polymer Refine Detection (Leica Biosystems) according to the manufacturer’s protocol. Anti–15-PGDH staining was performed using a commercially available antibody (Abcam, EPR14332-19, Catalog # ab187161). Whole-slide scanning (×40) was performed on an Aperio AT2 (Leica Biosystems).

### Quantification of cell populations

Flow cytometry was performed on a BD LSRII (BD Biosciences, San Jose, CA), and data were analyzed using FlowJo software (TreeStar). The following antibodies were used to quantify the populations below:

#### HSPCs

Cells were stained with antibodies to: CD3 (Biolegend, Catalog # 100206), Ly-6G/Ly-6C (Invitrogen, Catalog # 12-5931-82), B220 (Biolegend, Catalog # 103208), Ter119 (BD Biosciences, Catalog # 553673), CD11b (Biolegend, Catalog # 101208), Sca1+ (Invitrogen, Catalog # 11-5981-82), c-Kit+ (Biolegend, Catalog # 105812), CD48 (Biolegend, Catalog # 103422) and CD150 (Biolegend, Catalog # 115914).

#### Myeloid and Lymphoid

Cells were stained with antibodies to: CD11b (Biolegend, Catalog # 101212), CD3 (Biolegend, Catalog # 100204), B220 (Biolegend, Catalog # 103208), CD4 (Biolegend, Catalog # 116008), CD8 (Biolegend, Catalog # 100732), Ly6C (Biolegend # 128026), CD45 (Invitrogen # MCD4517), Ly6G (Invitrogen # 56-9668-82), and Live/Dead (Invitrogen # L34966A).

### Serum collection and inflammatory cytokine analysis

At least 400uL of blood was collected into BD Microtainer tubes (Beckton Dickinson Catalog # 365967) through submandibular cheek puncture. One hour after collection, blood was spun at 9000 rpm for 3 minutes, and at least 75uL of serum was aliquoted and frozen in Eppendorf tubes. Luminex® xMAP® technology was used to quantitatively and simultaneously detect thirty-two mouse cytokines (32-Plex Discovery Assay Array (MD32)), chemokines and growth factors. The multiplexing analysis was performed by Eve Technologies Corporation (Calgary, Alberta, Canada) using the Luminex® 200™ system (Luminex Corporation/DiaSorin, Saluggia, Italy) with Bio-Plex Manager™ software (Bio-Rad Laboratories Inc., Hercules, California, USA).

### Serial competitive transplants

Recipient C57/Bl6 mice were exposed to 10 Gy total body irradiation from a cesium source. Sixteen hours later, mice received a retro-orbital cell infusion of a 1:1 mixture of 1 × 10^6^ whole BM cells containing either 60 day Vehicle BM:naïve CD45.1 expressing B6.SJL-*Ptprc*^*a*^ *Pepc*^*b*^/BoyJ BM, or 60 day PGDHi BM:naïve CD45.1 BM. Every 2 weeks peripheral blood was collected from the submandibular vein to determine CD 45.2 frequency following competitive transplant (FITC Invitrogen, Catalog # 11-0454-81), PacBlue CD45.1 (Biolegend, Catalog # 110722), PE B220 (Biolegend, Catalog # 103208), PE CD3 (Biolegend, Catalog # 100206), APC CD11b (Biolegend, Catalog # 101212). At 16 weeks following primary transplant, chimeric mice were sacrificed, and BM was pooled from each experimental arm, and subsequently transplanted into another cohort of lethally irradiated recipient mice. Secondary transplant recipients were assessed for peripheral chimerism every 2 weeks for 12 weeks, after which animals were sacrificed, and BM composition was characterized via cytometric analysis.

### Statistics

All values were tabulated graphically with error bars corresponding to standard error of the means. Analysis was performed using GraphPad Prism software. Unpaired two-tailed Student’s t-test was used to compare groups, unless otherwise noted. For peripheral blood recovery kinetic analysis, 2-way ANOVA was used to test the effect of drug treatment. P<0.05 was considered statistically significant.

## Results

### 15-PGDH Expression and Activity are Conserved in the Aging Hematopoietic System

To assess age-related changes in 15-PGDH expression and localization, we analyzed bone marrow (BM) and spleens from young (2–3 months) and aged (18-month-old) mice. Aged animals exhibited a characteristic hematopoietic aging phenotype, including a myeloid-biased differentiation pattern, increased phenotypic HSPCs, and elevated pro-inflammatory signaling^28,29^ (**Supplemental Figure 1**). Immunohistochemical analysis revealed no differences in 15-PGDH protein localization between young and aged cohorts (**Figure 1A**). Similarly, *Hpgd* gene expression remained stable across age groups, though expression was significantly higher in the spleen compared to BM, as we previously reported ^30^ (**Figure 1B**). This indicates that both protein and gene expression of 15-PGDH are maintained throughout aging. Given that prostaglandin E2 (PGE2) signaling is mediated by the EP2 (*Ptger2*) and EP4 (*Ptger4*) receptors, we also examined their expression but found no significant differences between young and aged mice^31,32^ (**Supplemental Figure 2**). To determine whether enzymatic activity changes with age, we quantified 15-PGDH activity in bulk, CD11b+, F4/80+, and B220/CD3+ cell populations from BM and spleen. Consistent with previous findings in young mice, where 15-PGDH activity is enriched in F4/80+ macrophages^33^, this pattern was also observed in aged BM and spleen (**Figure 1C-D**). However, we detected a global reduction in enzymatic activity across all hematopoietic populations with age, suggesting that while 15-PGDH expression remains stable, its activity is modulated over time in a tissue-specific manner. To confirm that PGDHi effectively inhibits enzyme activity, we assessed bulk BM and splenic activity 30 minutes following a single injection of 10 mg/kg PGDHi. Treatment resulted in a significant reduction in 15-PGDH activity in both BM and spleen, confirming effective enzymatic inhibition in aged mice (**Figure 1E**).

**Figure 1:**
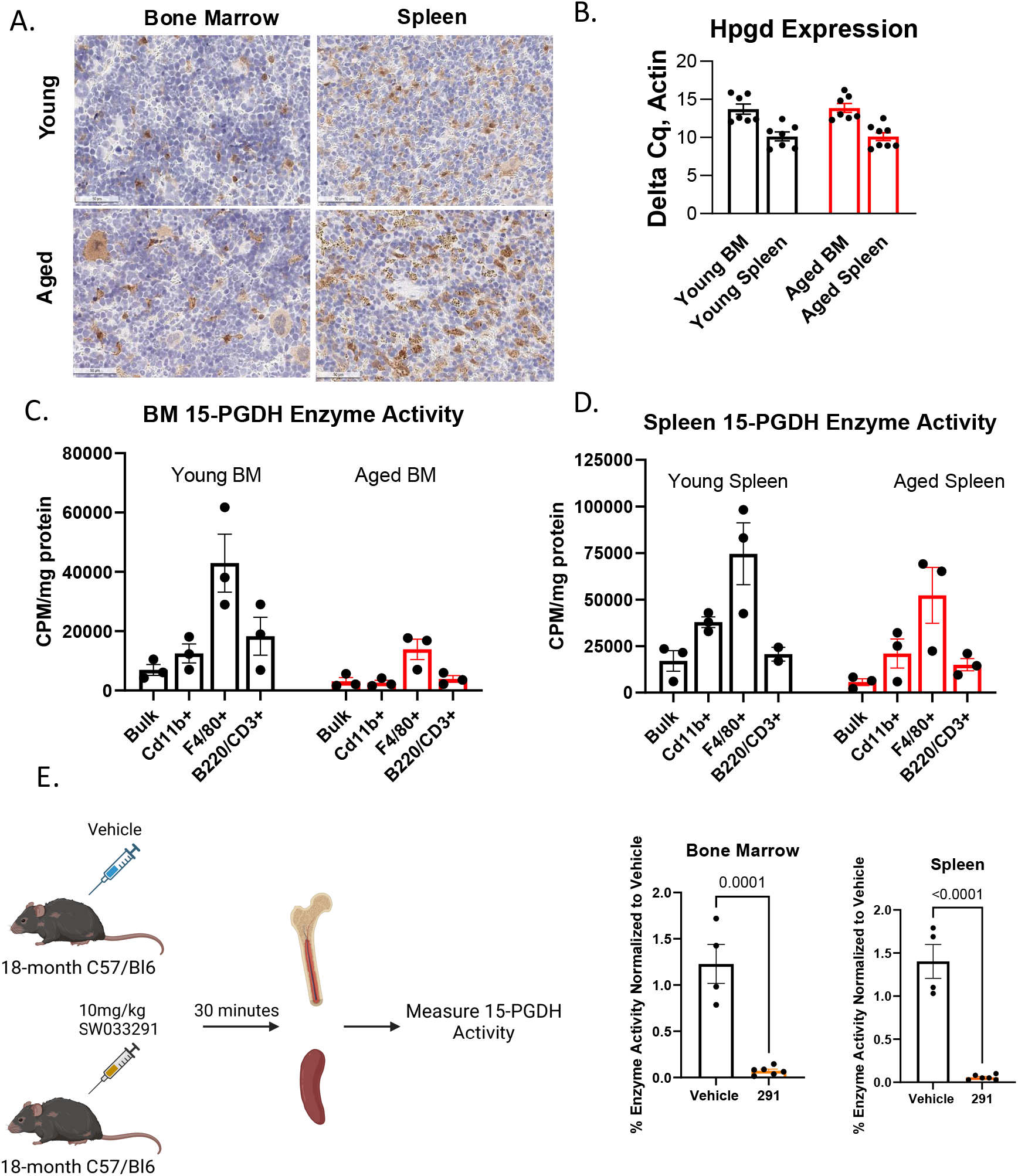
15-PGDH Expression and Activity are Conserved in the Aging Hematopoietic System. **A**. Representative images of 15-PGDH staining (brown) in naïve murine bone marrow, and spleen in young (2-3 months) vs. aged (>18 month) old mice. **B**. Relative gene expression of *Hpgd* in murine bone marrow and spleen by RT-PCR. Values were normalized to *Actb* levels and expressed as fold change relative to the level in young mice. N=7-8 mice/arm. **C, D**. Young and aged BM and splenic 15-PGDH activity in bulk, CD11b+, F4/80+, and B220/CD3+ purified populations expressed as counts per minute per mg of protein. **E**. BM and splenic 15-PGDH activity in aged mice 30 minutes following vehicle or 10 mg/kg SW033291 (PGDHi). N=4-5 mice/arm. Schematic made using BioRender.com.

**Figure 2:**
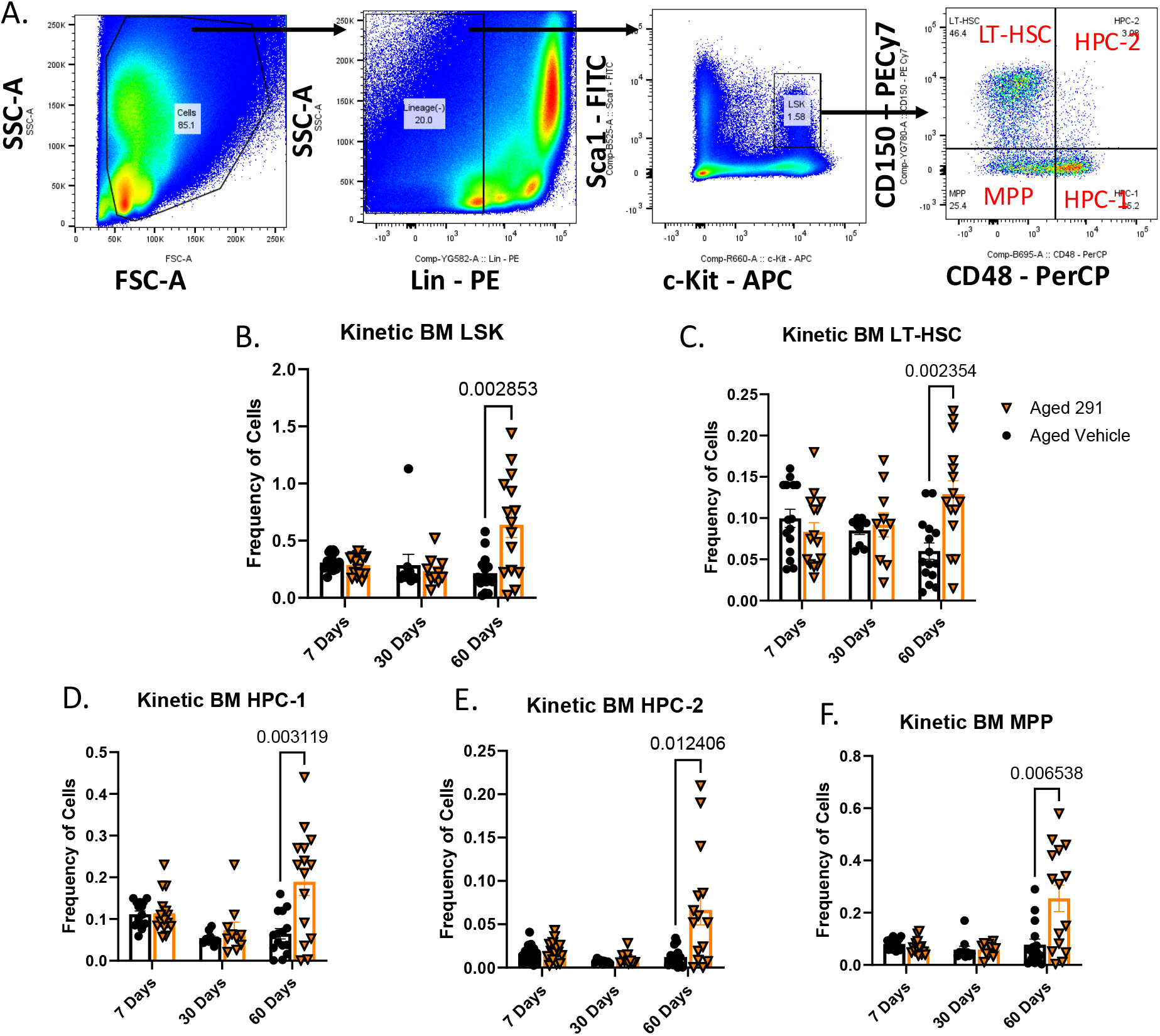
Sustained PGDHi Expands Bone Marrow HSPCs in Aged Mice. **A**. Representative flow cytometry dot plots of lineage(-)/Sca1(+)/c-Kit(+) (LSK), LSK/CD150(+)/CD48(-) (LT-HSC), LSK/CD150(+)/CD48(+) (HPC-2), LSK/CD150(-)/CD48(+) (HPC-1), and LSK/CD150(-)/CD48(-) (MPP) populations. **B-F**. Bone marrow LSK, LT-HSC, HPC-1, HPC-2, and MPP frequencies after 7, 30, and 60 days of treatment with vehicle or SW033291.

### Sustained PGDHi Expands Bone Marrow HSPCs in Aged Mice

Given that 15-PGDH expression and activity remain stable with age (**Figure 1**), we hypothesized that prolonged PGDHi treatment could counteract age-associated hematopoietic decline by expanding HSPC populations. To test this, aged mice were treated with PGDHi twice daily for 7, 30, and 60 days, followed by flow cytometric analysis of bone marrow HSPCs. Representative flow cytometry plots illustrate increased HSPC frequencies including lineage(-)/Sca1(+)/c-Kit(+) (LSK) cells, long-term HSCs (LT-HSCs; LSK/CD48(-)/CD150(+)), and progenitor subsets (HPC-1, HPC-2, and MPPs) (**Figure 2A**). While transient HSPC expansion was observed at early timepoints, PGDHi elicited robust increases in LSK and LT-HSC frequencies by day 60 (**Figure 2B-C**). Additionally, HPC-1, HPC-2, and MPP populations ^34^ were all significantly expanded, suggesting a broad effect on progenitor cell output at D60 (**Figure 2D-F**). A parallel systemic response was evident in the spleen, where HSPC populations were similarly elevated at day 60 (**Supplemental Figure 3A**).

### PGDHi Does Not Disrupt Steady-State Hematopoiesis In Aged Mice

To evaluate the lineage specificity of PGDHi, we analyzed mature lymphoid and myeloid populations in the bone marrow and peripheral blood of aged mice treated with PGDHi or vehicle for up to 60 days. Representative flow cytometry plots illustrate the gating strategy used to identify B220+ B cells, CD4+ and CD8+ T cells, and myeloid subsets in the BM and PB (**Figure 3A**). Quantification of BM populations demonstrated that total lymphoid cells, B cells, and CD4+/CD8+ T cells remained unchanged across all time points (**Figure 3B**). BM myeloid populations, including Ly6C-high and Ly6C-low subsets, were unaffected by PGDHi treatment (**Figure 3C**). Splenic lymphoid and myeloid lineages similarly remained unchanged with 30 days of PGDHi (**Supplemental Figure 3B-C**). Peripheral blood analysis further confirmed that PGDHi does not alter steady-state hematopoiesis. Myeloid, B cell, and T cell frequencies remained stable throughout the 60-day treatment period, while a modest decline in circulating myeloid cells was observed in PGDHi-treated mice (**Figure 3D**). Though this reduction was not statistically significant, it may suggest a shift away from age-associated myeloid skewing. Together, these findings indicate that PGDHi selectively expands the HSPC compartment while maintaining steady-state mature hematopoietic populations in aged mice, a key therapeutic advantage for mitigating age-related dysfunction without compromising blood homeostasis.

**Figure 3:**
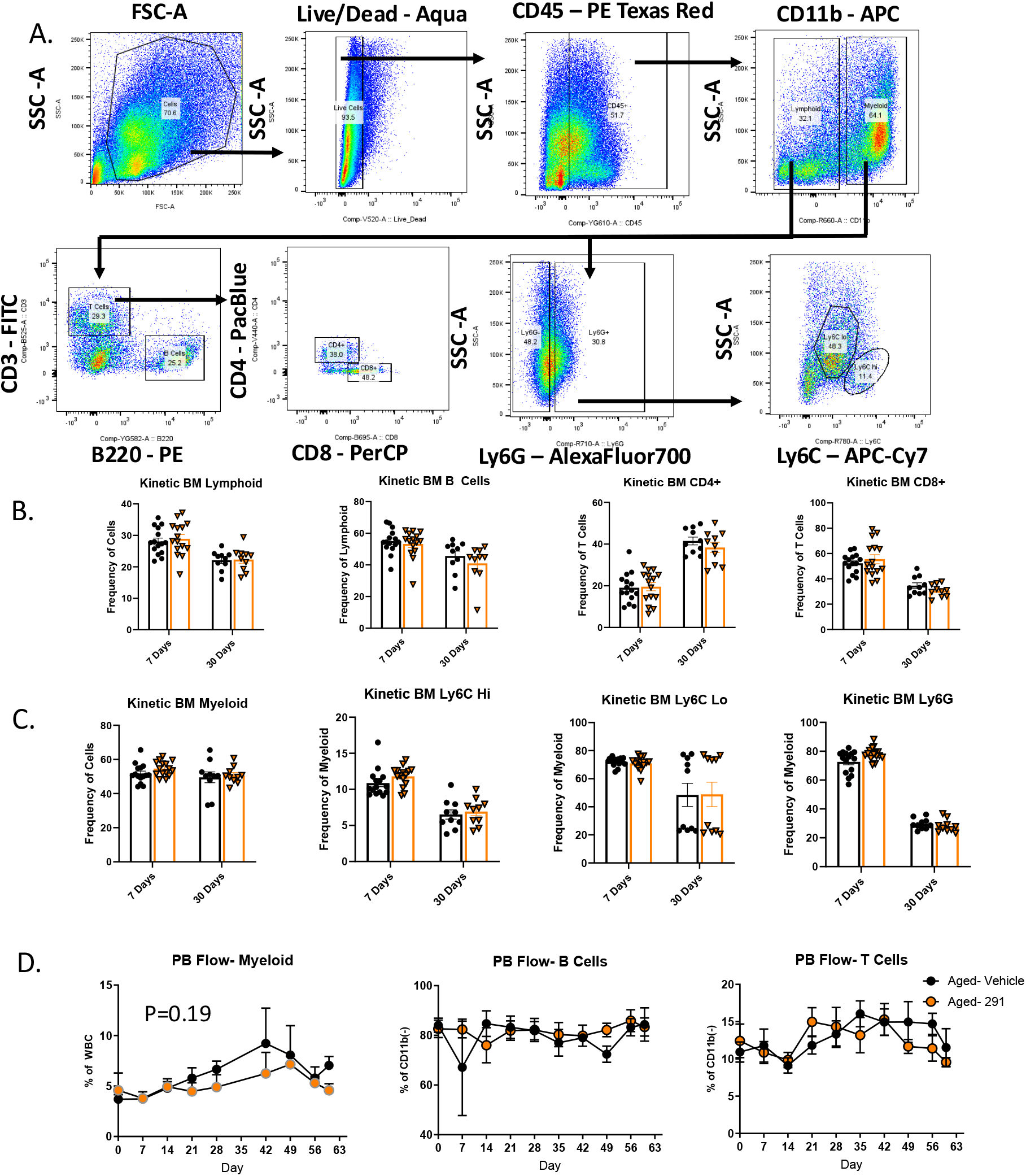
PGDHi Does Not Disrupt Steady-State Hematopoiesis In Aged Mice. **A**. Representative flow cytometry dot plots of lymphoid, B220+, CD4+, CD8+, myeloid, Ly6G, Ly6C Low, and Ly6C High cell populations from BM and spleen. **B**. Bone marrow total lymphoid, B220+ B Cell, CD4+ T Cell, and CD8+ T Cell populations from 7 (N=15) and 30 (N=10) days of twice daily vehicle or SW033291 treatment. **C**. Bone marrow total myeloid, Ly6C High, Ly6C Low, and Ly6G populations from 7 (N=15) and 30 (N=10) days of twice daily vehicle or SW033291 treatment. **D**. Peripheral blood flow cytometric analysis of myeloid, B cell, and T cell over 60 days of twice daily treatment with vehicle or SW033291. N=5 mice/arm.

### PGDHi Induces A Regenerative Signature In Aged Bone Marrow

To define the molecular mechanisms underlying PGDHi’s effects, we analyzed cytokine dynamics and transcriptional profiles in aged BM. Multiplex cytokine analysis from D30 serum revealed shifts in bone marrow cytokine expression following PGDHi treatment, with several factors trending toward decreased levels. IL-3 was significantly reduced, while IL-10, KC, RANTES, IL-12p70, MIP-2, and M-CSF all showed decreasing trends that did not hit significance (see **Supplemental Figure 4** for additional factors measured). IL-3 plays a role in hematopoietic progenitor proliferation and myeloid differentiation^35^, and its reduction may indicate a shift away from inflammatory myelopoiesis. Similarly, the trend in decreased IL-12p70 and MIP-2, which contribute to pro-inflammatory signaling and neutrophil recruitment^36,37^, suggest a suppression of inflammatory signaling. M-CSF, a regulator of monocyte and macrophage differentiation^38^, also exhibited a decreasing trend, suggesting a shift in cytokine signaling that may influence myeloid lineage potential under regenerative conditions, rather than steady-state myeloid composition. In contrast, LIF levels increased at day 30, a cytokine known to support hematopoietic stem and progenitor cell maintenance and promote self-renewal under stress conditions^39^ (**Supplemental Figure 5A**). Transcriptional profiling further revealed a pro-regenerative signature characterized by anti-inflammatory reprogramming and BM niche remodeling. With respect to anti-inflammatory reprogramming, we observed upregulation of *Arg1, Mrc1*, and *Tgf*β*1*, markers of tissue-reparative and anti-inflammatory consistent with M2 macrophage polarization. With respect to niche remodeling, we observed induction of HSC-supportive factors *Csf1, Kitl, Angpt1*, and *Spp1* after 60 days of PGDHi (**Supplemental Figure 5B and Supplemental Figure 6**). These findings demonstrate that PGDHi restores balance in the aged BM milieu, dampening inflammatory signals while activating programs essential for tissue repair and niche-mediated HSC support.

### PGDHi-Treated Donor Cells Accelerate Post-Transplant Recovery in Aged Models

To assess the therapeutic potential of PGDHi in hematopoietic stem cell transplantation (HSCT), we transplanted BM cells from aged donor mice pretreated with 30 days of PGDHi or vehicle into lethally irradiated recipients. PB and BM were analyzed 21 days post-transplant (**Figure 4A**). Recipients of PGDHi-treated BM exhibited significantly elevated white blood cells in PB, driven by a robust increase in neutrophils and a trend toward higher lymphocytes, suggesting enhanced hematopoietic reconstitution in the periphery following transplantation. Erythroid and platelet lineages remained unaffected, indicating lineage-specific enhancement (**Figure 4B-C**). BM analysis demonstrated a significant increase in total cellularity, while LSK and LT-HSC frequencies were reduced in both BM and spleen (**Figure 4D and Supplemental Figure 3D**), which may suggest accelerated mobilization of HSPCs from reservoirs to peripheral sites. These findings demonstrate that PGDHi preconditioning primes aged donor cells for rapid engraftment, promoting neutrophil-driven recovery while preserving erythroid / thrombopoietic homeostasis. The concurrent depletion of quiescent HSPCs in BM and spleen underscores a mechanism of enhanced differentiation and mobilization, critical for mitigating prolonged cytopenias in aged transplantation.

**Figure 4:**
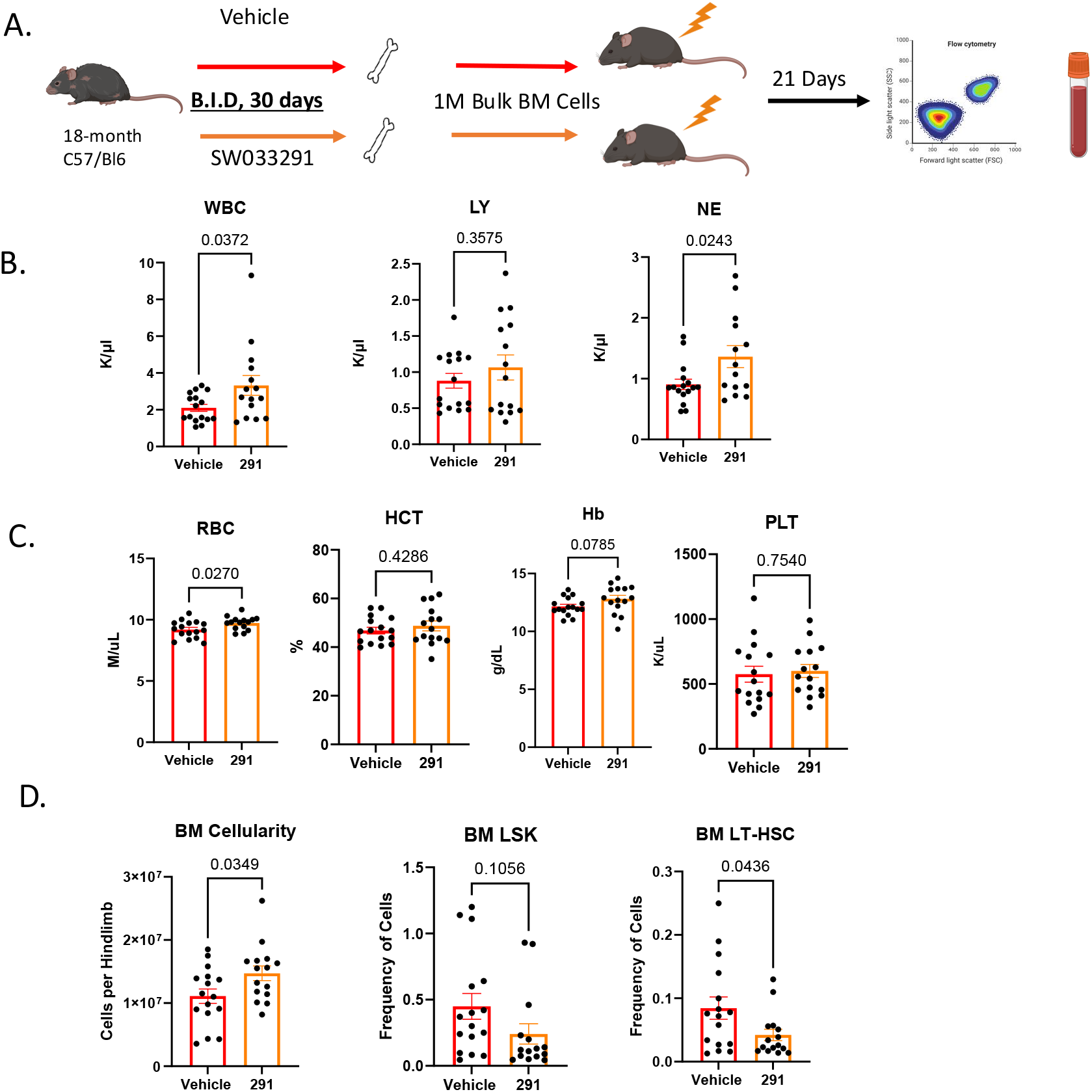
PGDHi-Treated Donor Cells Accelerate Post-Transplant Recovery in Aged Models. **A**. Schematic depicting experiment. 18-month C57/Bl6 mice were treated twice daily with vehicle or SW033291 for 30 days. 1 million bulk BM cells were transplanted into lethally irradiated 8-10 wk old mice. After 21 days, cellularity, flow cytometric data, and peripheral complete blood counts were acquired. **B-C**. Complete blood counts on peripheral white blood cells (WBC), lymphocytes (LY), neutrophils (NE), monocytes (MO), red blood cells (RBC), hematocrit (HCT), hemoglobin (Hb), and platelets (PLT). **D**. BM cellularity from one hindlimb; flow cytometric assessment of BM LSK and LT-HSC frequencies. N=15-16 mice/arm, 2 biological replicates. Schematic made using BioRender.com.

### PGDHi-Treated Aged BM Demonstrates A Competitive Advantage In Primary HSCT And Suppresses Myeloid Skew In Secondary HSCT

To assess the functional properties of aged HSCs following PGDHi treatment, we performed competitive transplantation studies using bone marrow from aged mice treated for 60 days with either PGDHi or vehicle. 5^e5^ bone marrow cells from these mice (CD45.2) were mixed 1:1 with young, congenic CD45.1-expressing BM and transplanted into lethally irradiated young recipients (**Figure 5A**). Peripheral blood chimerism was measured every two weeks post-transplant to track donor cell contribution. In primary transplantation, recipients of PGDHi-treated donor cells exhibited a significant competitive advantage over those receiving vehicle-treated BM, with higher CD45.2 chimerism in the peripheral blood over 16 weeks (**Figure 5B**). However, by the time of bone marrow collection at week 16, total donor chimerism in the BM was similar between groups (**Figure 5C**), suggesting that the early advantage observed in circulation was due to increased proliferative output rather than differences in long-term HSC engraftment. To evaluate how PGDHi-treated HSCs respond to additional replicative stress, we performed secondary transplantations by pooling BM from each group of primary recipients (Aged Vehicle:Naïve and Aged PGDHi:Naïve) and transplanting into lethally irradiated young recipients (**Figure 5D**). Over 12 weeks, peripheral blood chimerism was lower in recipients of PGDHi-treated donor cells compared to those that received vehicle-treated donor cells (**Figure 5E**). This difference was driven by a pronounced myeloid skew in the Vehicle group, with vehicle-derived donor cells comprising ∼90% of myeloid cells in competition with the naïve counterpart at Week 12 (**Figure 5F**). Since myeloid skew is a well-established feature of hematopoietic aging, these findings indicate that PGDHi treatment of aged HSCs reduces this bias, leading to a more balanced leukocyte output following transplantation.

**Figure 5:**
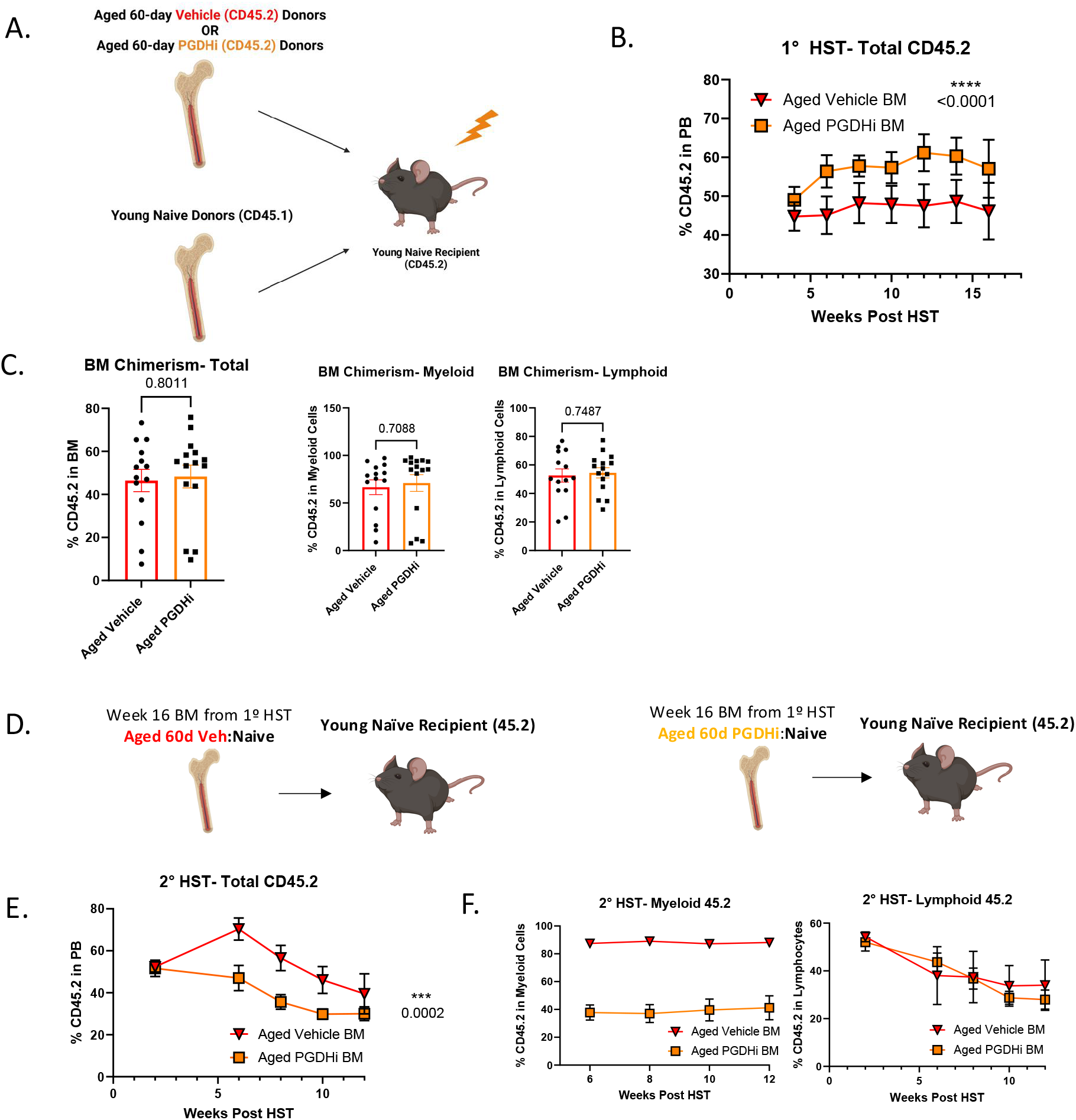
PGDHi-Treated Aged BM Demonstrates A Competitive Advantage In Primary HSCT And Suppresses Myeloid Skew In Secondary HSCT. **A**. Schematic depicting primary hematopoietic stem cell transplantation (HSCT). 500k bulk BM cells from aged (18 -month-old) mice treated for 60 days with vehicle or SW033291 (CD45.2 genotype) were transplanted alongside 500k bulk BM cells from young naïve mice (CD45.1 genotype) into lethally irradiated young recipients. **B**. 16-week peripheral blood recovery of mice that received vehicle or SW033291-treated CD45.2 donor cells. **C**. Total, myeloid-specific, and lymphoid-specific BM chimerism. N=15 mice/arm over 2 replicates. **D**. 12-week peripheral blood chimerism from secondary HST. **E**. Myeloid- and lymphoid-specific peripheral blood chimerism from secondary HST. N=5 mice/arm. Schematic made using BioRender.com.

### PGDHi Exhibits Favorable Safety Profile in Aged Mice

To evaluate the tolerability of prolonged PGDHi administration in aged mice, we analyzed complete blood counts (CBC) and serum chemistry profiles after 30 days of treatment. PB analysis revealed no significant alterations in WBCs, lymphocytes, monocytes, granulocytes, RBCs, hematocrit, hemoglobin, or platelet counts between PGDHi- and vehicle-treated cohorts (**Figure 6A**). Serum chemistry profiles further indicated no group-specific differences. Key markers of hepatic function, including ALT, AST, and alkaline phosphatase, and renal function (blood urea nitrogen) remained within comparable ranges (**Figure 6B**). These findings collectively demonstrate that chronic PGDHi is well-tolerated in aged-mice, with no evidence of hematological, hepatic, or renal toxicity.

**Figure 6:**
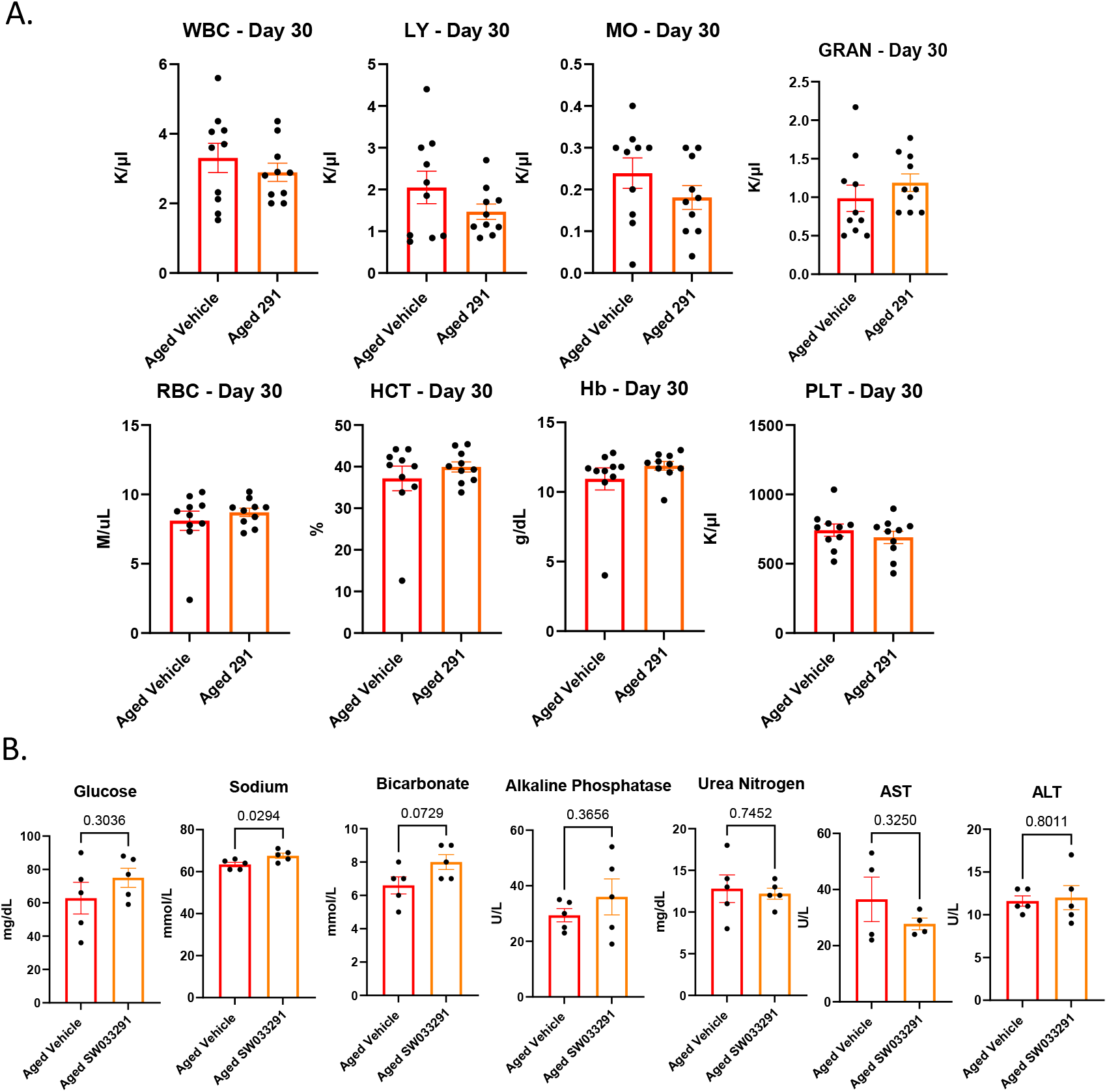
PGDHi Exhibits Favorable Safety Profile in Aged Mice. **A**. Peripheral complete blood counts on aged (>18mo) mice dosed twice daily for 30 days with either vehicle or SW033291. White blood cells (WBC), lymphocytes (LY), monocytes (MO), granulocytes (GRAN), red blood cells (RBC), hematocrit (HCT), hemoglobin (Hb), and platelets (PLT) were analyzed. N=10 mice/arm. **B**. Comprehensive serum chemistry on aged mice dosed twice daily for 30 days with either vehicle or SW033291. N=5 mice/arm.

## Discussion and Conclusions

Hematopoietic aging is characterized by diminished regenerative capacity, myeloid lineage bias, and elevated risk of hematologic dysfunction. Here, we show that PGDHi counteracts these age-related declines by expanding functional HSC pools, accelerating PB recovery post-transplantation, and restoring balanced lineage output, all without perturbing steady-state hematopoiesis or inducing systemic toxicity. Notably, aged HSCs from PGDHi-treated mice exhibited superior engraftment in primary transplants, suggesting enhanced reconstitution through improved proliferation, differentiation, or niche adaptation. In secondary transplants, vehicle-treated HSCs recapitulated the age-associated myeloid-skewing, while PGDHi-treated HSCs maintained balanced lineage commitment, indicating restoration of youthful regulatory networks.

These functional improvements align with PGDHi’s ability to reshape BM microenvironment into a pro-regenerative state. Reductions in inflammatory cytokines (IL-3, IL-12p70, and MIP-2) and upregulation of stromal maintenance factors (LIF, *Csf1*, and *Angpt1*) suggest attenuated inflammation and improved niche support. Given 15-PGDH’s enrichment in macrophages, its inhibition may further promote tissue-reparative immune polarization, as evidenced by signatures favoring tissue remodeling over pro-inflammatory signaling. Notably, PGE2 has been shown to promote M2 macrophage polarization^40^. This mechanistic interplay positions PGDHi as a dual modulator of both HSC function and niche dynamics.

Beyond its effects on HSC function, these findings have broader therapeutic implications. Chronic inflammation is increasingly recognized as a driver of clonal hematopoiesis of indeterminate potential ^41,42,43^, which is linked to increased risks of hematologic malignancies and cardiovascular disease. Since PGDHi reduces inflammatory signaling in aged bone marrow, future studies should assess whether it can suppress the expansion of CHIP-associated clones or modify their functional properties. Given that myeloid skewing is a precursor to hematologic disease, sustained PGDHi treatment could also be explored as a strategy to mitigate age-related hematopoietic dysfunction and reduce the risk of myeloid malignancies. Additionally, PGDHi’s ability to enhance early hematopoietic reconstitution suggests a potential role in improving outcomes for patients with poor graft function following hematopoietic stem cell transplantation ^44 45^. In chemotherapy-induced neutropenia, where rapid hematopoietic recovery is crucial to reducing infection risk and maintaining treatment schedules, PGDHi’s ability to enhance early reconstitution suggests it could improve post-chemotherapy hematopoietic recovery, especially in older patients with diminished reserves.

While our findings establish PGDHi as a compelling therapeutic target to enhance hematopoietic function during aging, key questions remain. Future work should investigate whether chronic PGDHi prevents HSC exhaustion beyond 60 days. The molecular drivers of PGDHi-induced HSC expansion and lineage rebalancing should also be assessed via omics-based approaches, including single-cell transcriptomics and proteomics. Additionally, while no toxicity was observed in aged mice, broader safety assessments in preclinical models, as well as studies in human HSCs, will be critical for clinical translation.

In conclusion, our work challenges the paradigm that HSC rejuvenation requires disrupting steady-state hematopoiesis. By selectively modulating the BM niche, PGDHi enhances regenerative capacity while preserving homeostatic function, representing a therapy with significant potential for aging populations and patients undergoing hematopoietic stress.

## Acknowledgements

We thank Dr. Joseph Ready, Professor at UT Southwestern Medical Center, for performing QC analysis and providing aliquots of (+)SW033291 for these studies, and also for his input on experimental design and directions. We also thank the Cytometry and Microscopy Shared Resource at Case Western Reserve University and the Cleveland Clinic Lerner Research Institute for their support in developing flow cytometry panels and providing imaging services for tissue sample analysis.

## Funding Sources

This work was supported by NIH grants R21 AG075573, R00 HL135740, RM1GM42002, R35 CA197442, by the Case Comprehensive Cancer Center Support Grant P30CA043703 (Radiation Resources Core Facility, the Hematopoietic Biorepository and Cellular Therapy Core Facility, and the Cytometry & Imaging Microscopy Core Facility). This project was also supported by the Clinical and Translational Science Collaborative of Northern Ohio which is funded by the National Center for Advancing Translational Sciences (NCATS) of the National Institutes of Health, UM1TR004528, and a grant from Ohio Cancer Research. A.A.P. was supported by The Valour Foundation, as the Rebecca E. Barchas, MD, University Professor in Translational Psychiatry of Case Western Reserve University, as the Morley-Mather Chair in Neuropsychiatry of University Hospitals of Cleveland Medical Center, and through the Louis Stokes VA Medical Center resources and facilities. Financial support for this work was also provided by the NIDDK Innovative Science Accelerator Program (ISAC, www.isac-kuh.org), grant DK128851.

## Conflict of Interest Disclosures

ABD and SDM, and AAP are inventors on patents describing inhibitors of 15-PGDH and their therapeutic uses. These patents have been licensed to Rodeo Therapeutics, which is owned by Amgen.

## Data Availability Statement

All data underlying this article are available in the article and in its online supplementary material.

## Notes

### Competing Interest Statement

The authors have declared no competing interest.

