## Supplemental Figures for "15-PGDH inhibition promotes hematopoietic recovery and enhances HSC function during aging"

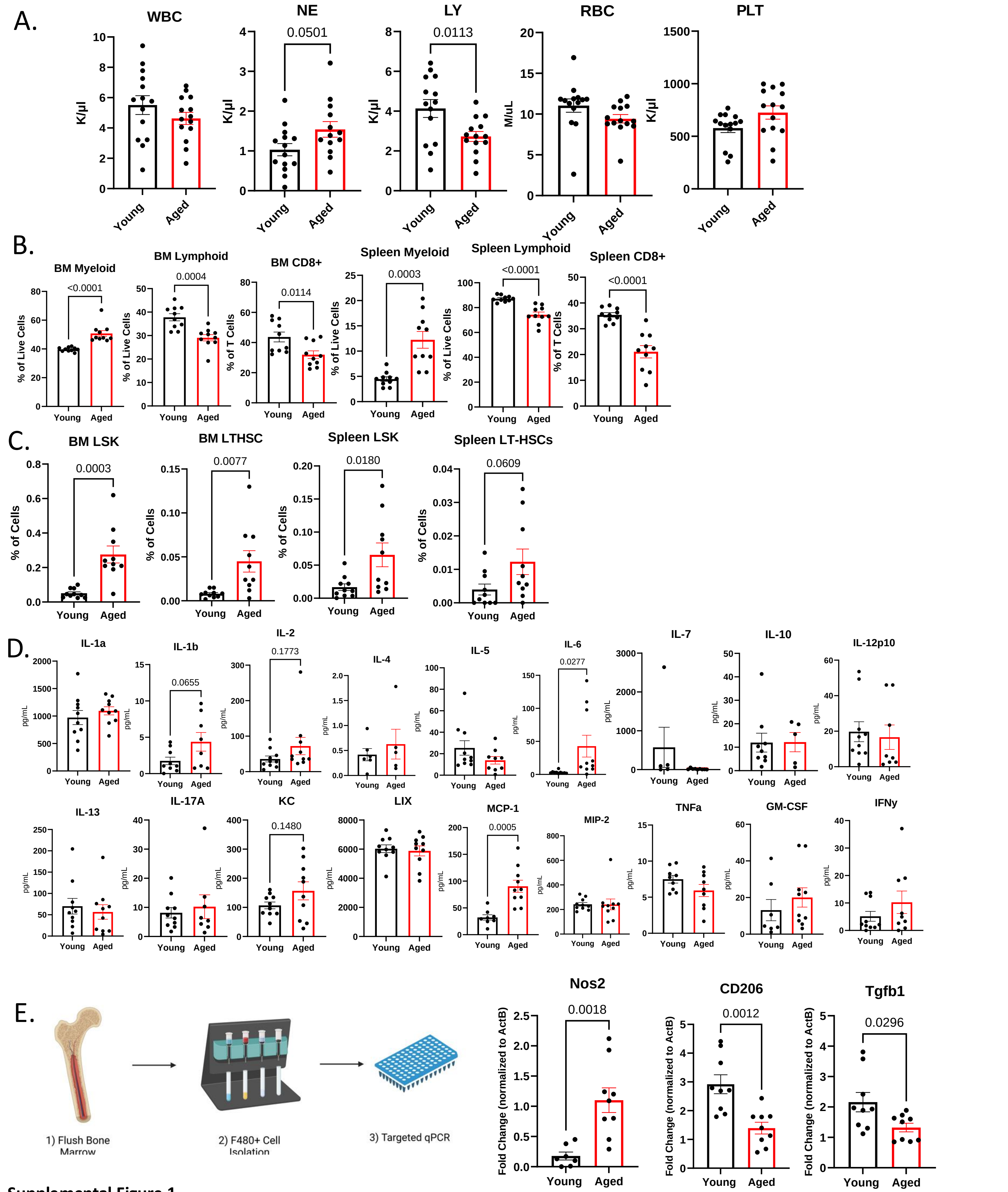


**Supplemental Figure 1**

**A.** Peripheral complete blood counts (CBCs) on naïve young (2-3 month) and aged (>18month) mice. N=14 mice/arm. **B.** Cytometric assessment of myeloid and lymphoid populations in bone marrow and spleen. N=10 mice/arm. **C.** Immunophenotypic analysis of hematopoietic stem and progenitor cells (HSPCs; Lineage- Sca1+ c-Kit+ (LSK)), and hematopoietic stem cells (HSCs; LSK CD48- CD150+ (SLAM)). N=10 mice/arm. **D.** Serum inflammatory cytokine quantification. N=10 mice/arm. **E.** Gene expression on purified F480+ bone marrow cells. N=10 mice/arm. Error bars represent SEM. Student's T-test performed for analyses. Schematic made using BioRender.com.

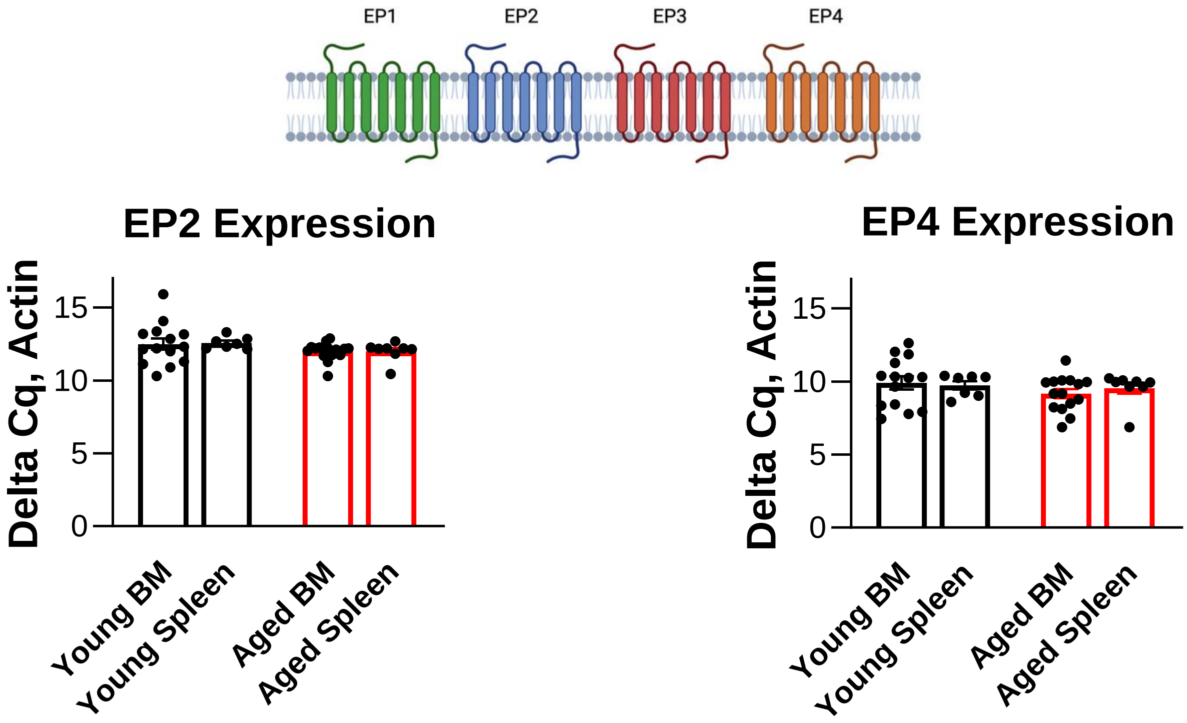


**Supplemental Figure 2**

**A.** Ptger2 (EP2) and Ptger4 (EP4) expression in young vs. aged mice. N=14-15 mice/arm for bone marrow and N=7-8 mice/arm for spleen.

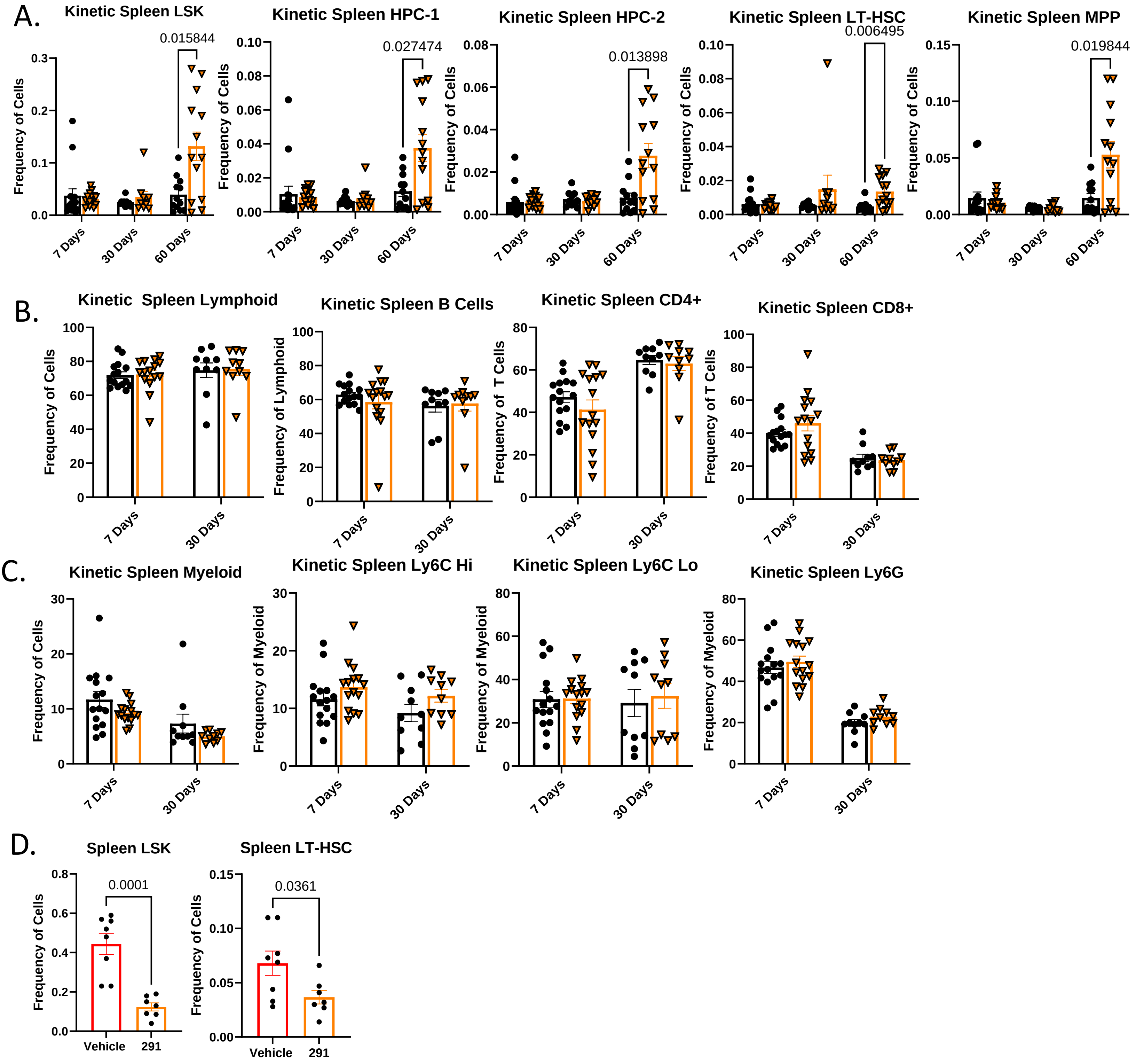


**Supplemental Figure 3**

**A.** Splenic LSK, LT-HSC, HPC-1, HPC-2, and MPP frequencies after 7, 30, and 60 days of twice daily treatment with vehicle or SW033291. **B.** Splenic total lymphoid, B220+ B Cell, CD4+ T Cell, and CD8+ T Cell populations from 7 (N=15) and 30 (N=10) days of twice daily vehicle or SW033291 treatment. **C.** Splenic total myeloid, Ly6C High, Ly6C Low, and Ly6G populations from 7 (N=15) and 30 (N=10) days of twice daily treatment. **D.** Splenic LSK and LT-HSC 21-days after short-term HST. 18-month C57/Bl6 mice were treated twice daily with vehicle or SW033291 for 30 days. 1 million bulk BM cells were transplanted into lethally irradiated 8-10 wk old mice. N=7-8 mice/arm.

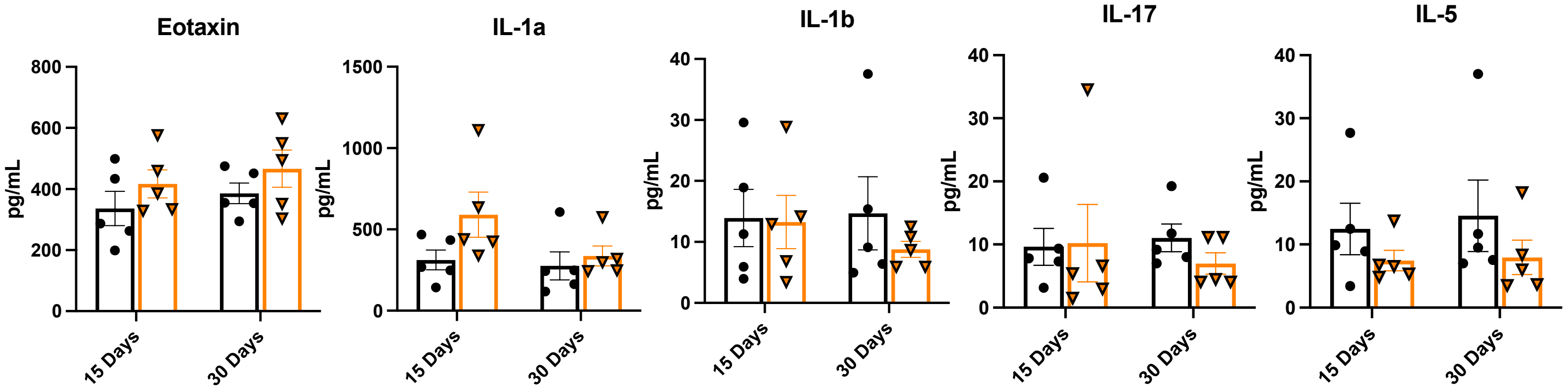


**Supplemental Figure 4**

Additional serum inflammatory analytes from 15 and 30 days of treatment with vehicle or SW033291. N=5 mice/arm.

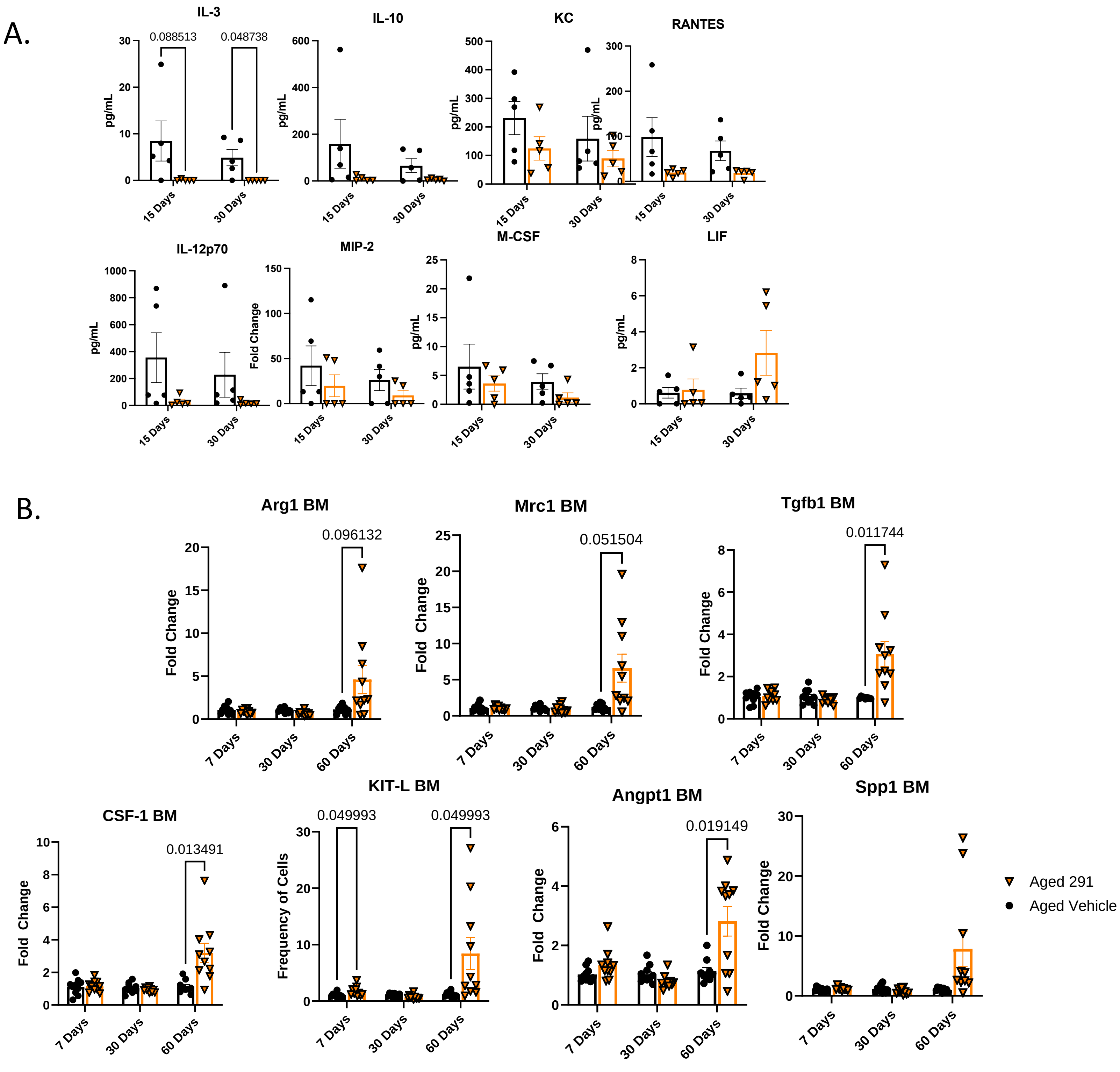


**Supplemental Figure 5: 15-PGDHi induces a regenerative signature in aged bone marrow**

**A.** Serum inflammatory analytes at 15 and 30 days of treatment. N=5 mice/arm. **B.** RT-PCR on macrophage and hematopoietic-relevant genes from bulk bone marrow at 7, 30, and 60 days of treatment. N=10-15 mice/arm.

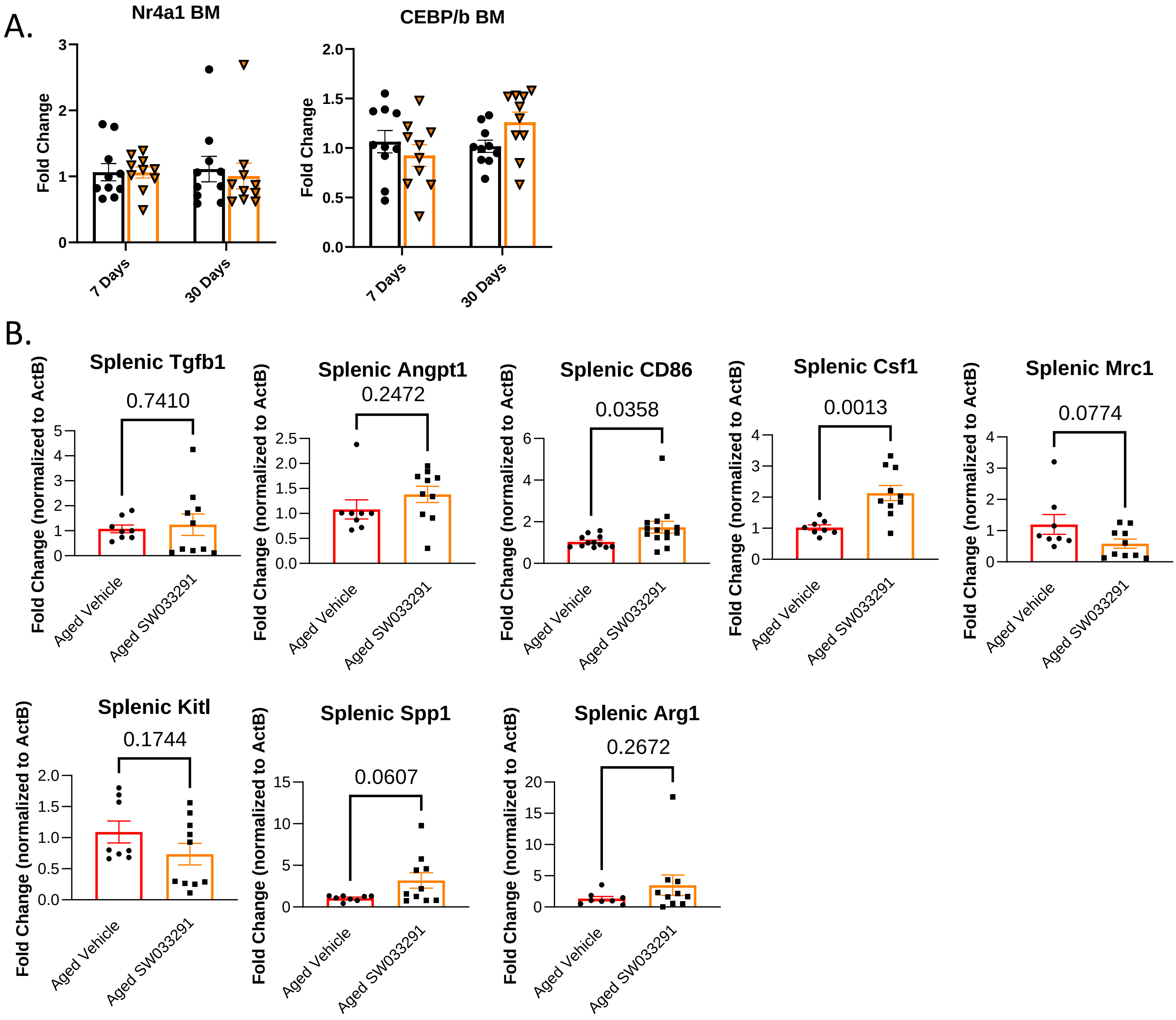


**Supplemental Figure 6**

**A.** Additional RT-PCR on macrophage-relevant genes from bulk bone marrow at 7 and 30 days of treatment with vehicle or SW033291. N=10 mice/arm. **B.** Splenic RT-PCR after 60 days of treatment. N=8-15 mice/arm
